# Integrating Career Development and Immigration Planning: A Pilot Program for International Scholars

**DOI:** 10.1101/2025.08.18.670737

**Authors:** Caroline Fecher, Natalie Chernets, Paola Cépeda

**Author notes:** N.C. was affiliated with Drexel University at the time of the study and is currently an independent consultant of workforce development. These authors contributed equally to this work (Natalie Chernets, Paola Cépeda). Corresponding author: Paola Cépeda.

## Abstract

International graduate students and postdoctoral researchers in the United States face unique challenges at the intersection of career development and immigration planning. Despite their high representation in U.S. research institutions, international scholars often lack access to integrated resources that address how their immigration status influences career mobility, decision-making, and long-term career planning. To address this gap, we developed the Effective Career and Immigration Planning for International Scholars (ECIP) program, a seven-session workshop series designed to align participants’ career development with their U.S. immigration goals. Grounded in Social Cognitive Career Theory, the ECIP program introduces the concept of the immigration portfolio: a strategic collection of evidence documenting accomplishments and credentials valuable for both academic success metrics and an advanced degree holder’s permanent residence petition. Here, we present the pilot implementation of this program and report its evaluation, which included pre- and post-program surveys, weekly polls of call-to-action task, and follow-up communications with participants.

Our results demonstrate significant increases in participants’ self-reported knowledge and confidence to take action towards their career and immigration goals. Post-program survey responses showed higher agreement levels on understanding immigration portfolio preparation and permanent residence options, demonstrating the program’s effectiveness in addressing knowledge gaps. Engagement data revealed that participants consistently completed the proposed action items from workshops, aligning with the session’s learning objectives. Participants considered the program relevant and empowering. Follow-up responses indicated continued progress toward the immigration goals of participants.

Our findings underscore the critical need for integrated career and immigration planning. The ECIP program offers a scalable, evidence-based model for supporting international scholars in navigating their professional development in the United States.

## 1. Introduction

Navigating academia as a graduate student or postdoctoral researcher is inherently demanding. Despite advanced training, early-career scholars face intense competition for limited resources such as funding, mentorship, and job opportunities (1,2). In response, many institutions in the United States have implemented structured career development programs to support scholars’ success (3–5). However, these frameworks often fail to account for the unique challenges faced by international scholars (6), who comprise 45% of graduate students and 60% of postdoctoral researchers in the United States (7–10). International scholars often encounter unspoken academic norms, language barriers, and cultural differences that delay integration into professional communities (11). Moreover, they must rebuild professional networks in the United States from the ground up, often without the social capital or institutional knowledge available to domestic peers (12,13). These factors significantly influence their career decision-making but are rarely addressed in existing career development programs (14).

A critical, underexplored dimension of career development is the impact of immigration status on career mobility (15) and long-term career opportunities in the United States (10). Temporary visa regulations affect access to internships, fellowships, and entrepreneurial ventures, which constitute key steppingstones for several career paths available to graduate degree recipients (16,17). Many international scholars, especially postdoctoral researchers, remain in lower-paid academic positions longer than they intended to maintain their legal status while pursuing sponsorship opportunities through prospective employers or applying for permanent residence (14,18,19) International graduate students in the United States often use the Optional Training Program (OPT) or other academic training programs associated with their visa to gain experience post-graduation and seek a work visa based on their track record with their employer. If unsuccessful, they may divert to postdoctoral researcher positions to maintain legal status. These challenges are further exacerbated by uncertainties around visa timelines, and the high cost of legal assistance.

Despite these barriers, a growing majority of international graduate students (73%) and postdoctoral researchers (>50%) expresses a desire to pursue long-term careers in the United States (20–22). As accomplished individuals in the sciences, arts, education, and other fields, graduate students and postdoctoral researchers could pursue employment-based permanent residence categories such as EB-1 or EB-2 (23). However, few resources exist to help them understand how their professional accomplishments can be used to strengthen their chances of both launching independent careers and obtaining permanent residence in the United States. The gap in resources causes uncertainty, diminishes confidence, and results in missed opportunities, which in turn increase stress and anxiety while decreasing productivity (24–27).

Additionally, institutional support services for graduate students and postdoctoral researchers frequently operate in isolation. On the one hand, career services are typically best equipped to guide scholars with domestic citizenship, as they frequently overlook visa restrictions that affect access to experiential opportunities. Career advisors may be unaware that pursuing non-traditional career paths can complicate the process of obtaining permanent residence. This is because it is harder to demonstrate the scholar’s future impact in a new field, requiring them to build new evidence to support their case and thus delaying their path to permanent residence. On the other hand, international offices tend to prioritize visa compliance over immigration education, often failing to connect immigration guidance with career planning. As a result, international scholars are left without support in navigating one of the most critical aspects of their professional journey: aligning career choices with long-term immigration goals (28,29).

Collectively, these factors underscore the intimate connection between career and immigration decisions. This study asserts that immigration decisions are not peripheral but central to the career development of international scholars (28). Here, we evaluated the pilot implementation of a novel professional development program, Effective Career and Immigration Planning for International Scholars (ECIP), which integrates immigration strategy into career planning. We tested the hypothesis that this integration enhances international scholars’ confidence, awareness, and self-efficacy in navigating their professional pathways in the United States. Moreover, we examined whether taking part in the ECIP program enabled participants to prioritize activities that simultaneously advance career goals and improve eligibility for permanent residence, thereby maximizing their long-term return on investment.

### The ECIP Program

The U.S. immigration system offers several employment-based pathways to permanent residence for highly trained individuals (23). Although categories like EB-1 for extraordinary ability closely mirror traditional academic success metrics (e.g., publications, peer review activity, and prestigious awards), these criteria are rarely addressed in academic training. We want to emphasize that this alignment may further enhance international scholars’ motivation to focus on scholarly productivity, an area in which they already outperform their domestic peers despite receiving lower compensation (18,30). This reinforcement highlights the dual value of such efforts for both academic advancement and immigration outcomes.

In prior work, we introduced the concept of the immigration portfolio (28) to underscore the parallels between immigration eligibility requirements and academic achievement metrics. An immigration portfolio is a curated, comprehensive collection of documents that demonstrate an individual’s qualifications and achievements in alignment with U.S. immigration criteria. It is analogous to a tenure dossier or faculty job application package, but it emphasizes intellectual contributions with the specific aim of establishing eligibility for permanent residence. For example, the EB-1 category requires evidence of extraordinary ability, including authorship of scholarly articles, experience judging the scholarly contributions of colleagues and peers, media recognition of expertise and professional achievements, and leadership roles in professional organizations. By highlighting the scholar’s unique expertise and potential to advance their field, the immigration portfolio can also support arguments for waiving employer sponsorship in the national interest (31). As the evidence collected mirrors the benchmarks of academic excellence, the immigration portfolio can serve as a strategic, dual-purpose tool for international scholars to align their professional development with long-term immigration goals.

Despite its importance, few international scholars receive training on how to build an immigration portfolio. Many are unaware of the need to document key activities, such as peer review contributions, media coverage of their work, or service as evaluators at scientific conferences. As a result, they often miss opportunities to strategically strengthen their immigration portfolio as they are not informed about the types of accomplishments that can demonstrate long-term value to the United States.

To address this gap, we designed the Effective Career and Immigration Planning for International Scholars (ECIP) program, which frames the development of the immigration portfolio as a strategic professional development practice. It provides international scholars with the knowledge, tools, and confidence to actively align their academic accomplishments with long-term career and immigration goals. Drawing on Social Cognitive Career Theory, which emphasizes self-efficacy and contextual influences on career development (32–34), the ECIP program aims to:

(1) Raise awareness of the intersection between career and immigration planning;
(2) Equip participants to strategically prioritize professional development activities that strengthen their immigration portfolio; and
(3) Foster peer support networks for accountability.

The ECIP program evolved from a postdoctoral peer support group founded by Dr. Natalie Chernets in 2016 at Drexel University. This group of international postdoctoral researchers met biweekly to exchange information on permanent residence petitions, review immigration categories, share cultural insights, expand professional networks, and peer-review materials. Meetings featured guest speakers and peer-led discussions, including reviews of successful petition examples that the members had compiled. While informal and lacking structured objectives or assessments, the group provided critical support for each other to address the professional challenges navigating immigration, academia, and life in the United States.

Building upon this foundational model, Dr. Natalie Chernets and Dr. Paola Cépeda introduced pedagogical enhancements beginning in 2020 to advocate for more inclusive career services with emphasis on the needs of international scholars. In 2022, they delivered a call-to-action presentation to career services professionals (35), piloted a 90-minute workshop at the University of Delaware (36), and published an opinion piece addressing strategies to support the career advancement of international scholars (28).

The workshop further evolved across multiple implementations, including a session for the National Postdoctoral Association’s Smart Skills Series in 2023 (37).

Developed with support from the Burroughs Wellcome Fund in 2023 (38), the current ECIP curriculum expands upon the previous 90-minute format by offering a multi-workshop program that integrates didactic content with applied activities and individually motivated work. It maintains the peer support model from the original 2016 group yet is strengthened by the incorporation of evidence-based practices in career development (32–34).

The ECIP program consists of seven 90-minute workshops, each designed to help participants master specific learning outcomes. These outcomes include understanding the intersection of career development with immigration education, exploring permanent residence pathways and associated challenges, organizing an immigration portfolio, applying effective reference letter strategies, developing professional CVs, and setting clear career-immigration goals (**Figure 1**). Importantly, the program does not provide legal advice but encourages informed exploration and education within the constraints of each participant’s immigration status.

**Figure 1.**
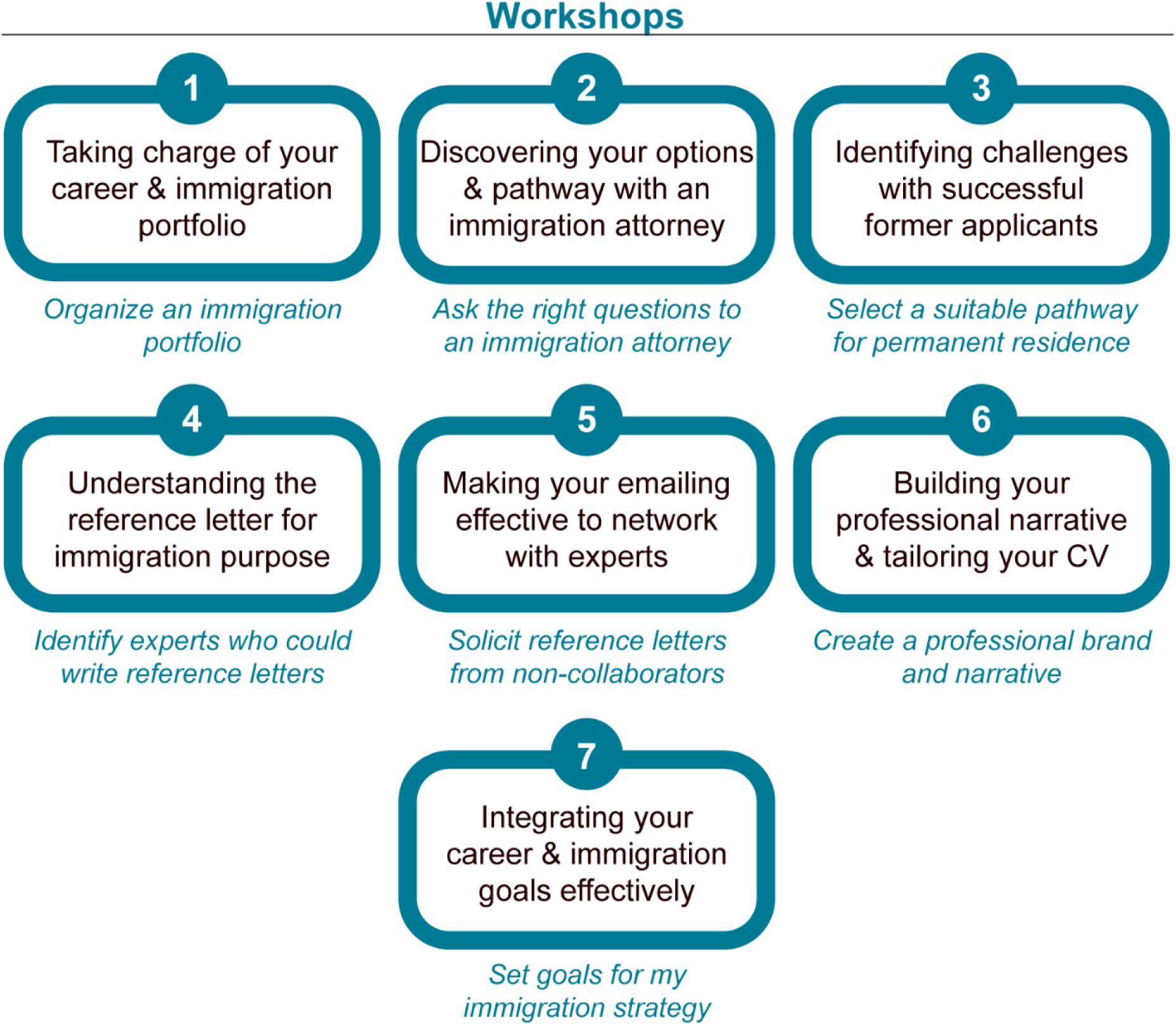
Effective Career and Immigration Planning for International Scholars (ECIP) Program Overview. The seven workshop topics (box) and their assigned learning outcomes (below box, blue).

In 2024, we piloted the ECIP program with 44 participants from two private R1 institutions in the United States: Drexel University in Philadelphia, PA, and Washington University in St. Louis, MO. Participants included postdoctoral researchers (52%), graduate students (41%), and research staff (7%). During this implementation (**Figure 2**), each workshop was organized into three sections. First, participants completed polls to report their progress on call-to-action tasks intended to develop their immigration portfolio and their resume. Second, participants engaged in an instructional component to learn the content and solidify their understanding using different approaches and learning strategies. Third, participants engaged in small cohort discussions designed to foster peer-to-peer support, enhance progress, and promote accountability. Peer cohorts were formed based on self-reported timelines for submitting a permanent residence petition, as reported in the pre-program survey. Specifically, participants were assigned to one of three interval groups: within 0-6 months, 6-12 months, or more than 12 months. The grouping was designed to facilitate targeted discussions and goal setting, aligning the workshop content with the participants’ specific stage in the permanent residence application process. These cohorts remained constant unless low workshop attendance necessitated the merging of formed groups.

**Figure 2.**
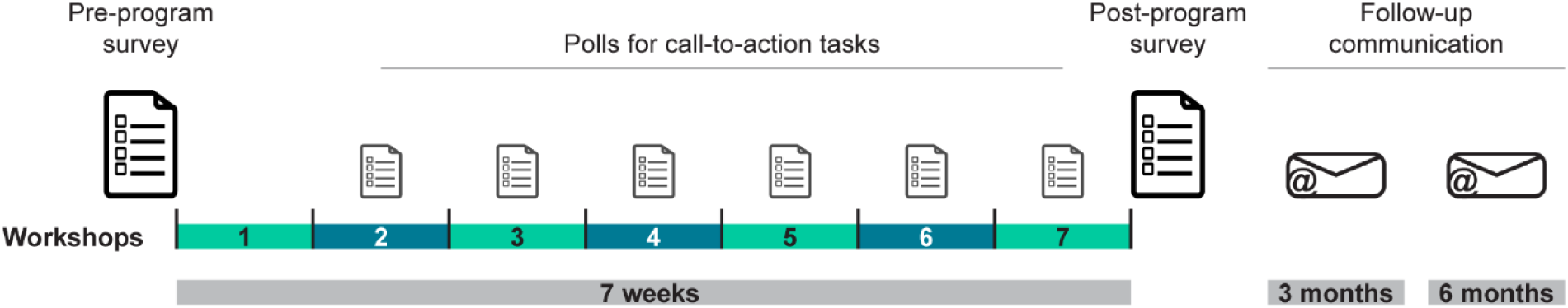
Evaluation of the ECIP program. Evaluation components across the ECIP program, containing pre- and post-workshop surveys, polls for call-to-action tasks in workshop #2-7, and follow-up communications after the program.

This article reports on the evaluation of this pilot implementation. We sought to compare participants’ self-reported career and immigration planning skill levels across all seven learning outcomes before and after the ECIP program. We also assessed perceived changes in knowledge and confidence to take action at the intersection of career and immigration decisions, as well as participants’ engagement and satisfaction with the program. The research questions that guided our analysis were:

1. How do participants self-assess their career and immigration planning skills across all seven learning outcomes?
2. How does the ECIP program influence participants’ knowledge and confidence in taking strategic actions to align their career and immigration goals?
3. What is the level of participant engagement with the workshop tasks?
4. How do participants evaluate the relevance and quality of the ECIP program for their professional development?

## 2. Materials and Methods

### 2.1. Study design

Evaluation of the program was built in at different time points: a pre-program survey, weekly polls inquiring about participants’ call-to-action tasks, a post-program survey, and follow-up communication after the program concluded with participants who completed the program (**Figure 2**). Participant consent was given by clicking a “Continue” button in the surveys. Data was either collected anonymously or de-identified after collection and aggregation. The study was submitted to the Institutional Review Board at Washington University in St. Louis under the ID #202308114 and considered as exempt from a full review.

The pre- and post-program surveys contained a core set of 31 identical questions for participant assessment and were administered before the first workshop and at the end of the last workshop, respectively. The questions asked participants to express agreement or disagreement on a 5-point Likert scale: 1-Strongly Disagree, 2-Disagree, 3-Neutral, 4-Agree, or 5-Strongly Agree. These questions addressed content from the seven workshops by asking about participants’ knowledge ("Knowledge") of the program topics and their confidence in taking related actions ("Action"). Participants were also asked to assess their perceived success in meeting the seven workshop learning outcomes (**Figure 1**). The pre-program survey included questions related to participant demographics, and the post-program survey contained questions about participants’ satisfaction with the program. Both surveys were conducted anonymously, and participants created a unique identifier to link their pre- and post-program responses.

Each workshop ended with call-to-action tasks for participants to complete by the following week. These tasks provided guided instructions aimed at developing and organizing their immigration portfolio. During the first part of workshops #2-7, participants completed polls to report on whether these tasks had been completed. Responses were tracked and accumulated across all polls to capture participants’ engagement and motivation regarding their career and immigration journeys.

Finally, follow-up communications were sent via email to participants who completed the program after three and six months. This communication asked participants about their progress regarding their self-set goals from workshop #7 and whether they had any additional information they wished to share.

### 2.2. Participants

Study participants are those who completed either the pre- or the post-program survey or both (**Table 1**). The total number of study participants was 44, all of whom completed the pre-program survey. Only 12 of these participants completed the post-program survey, with 11 respondents being matched to their pre-program survey identifier.

**Table 1.**
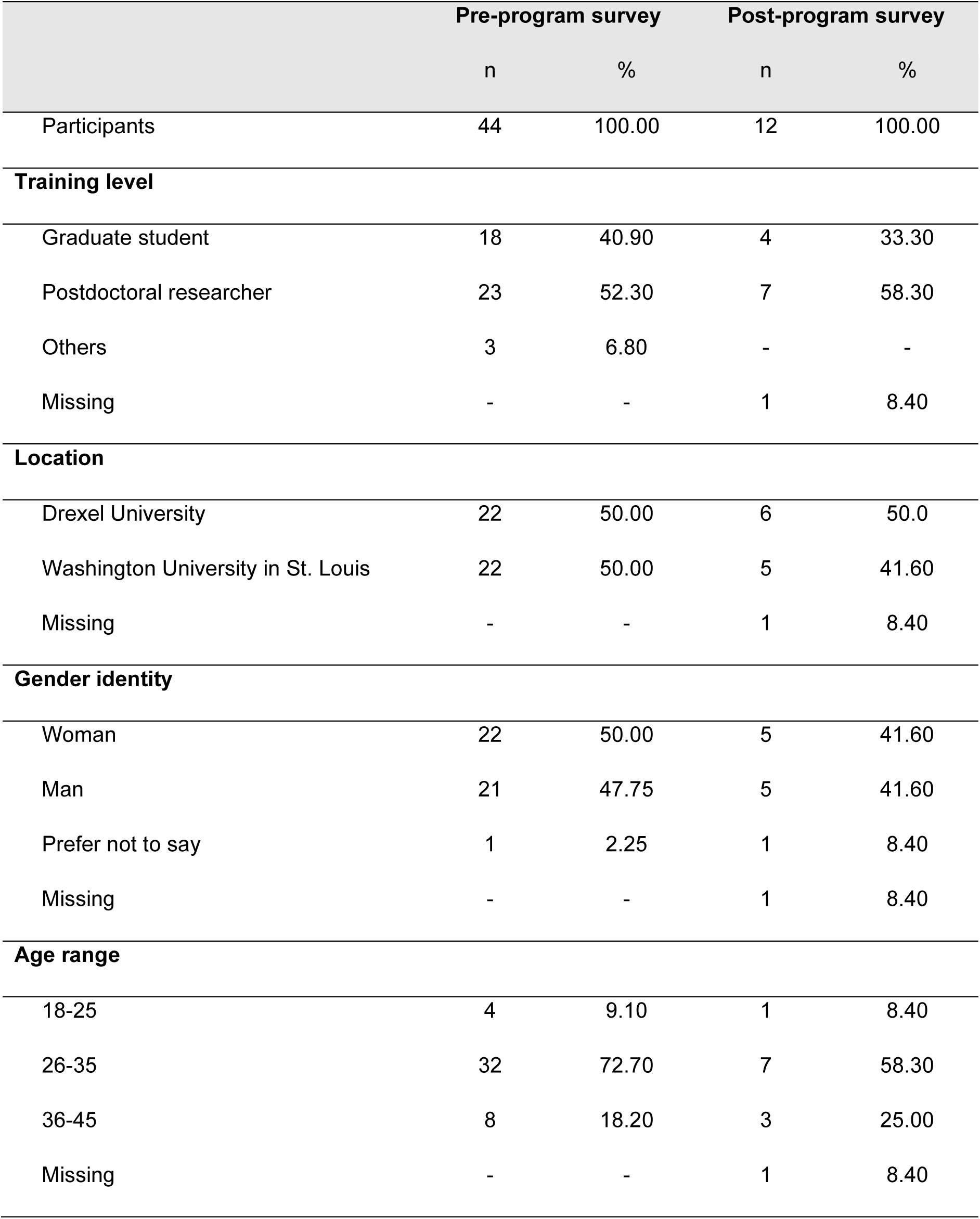
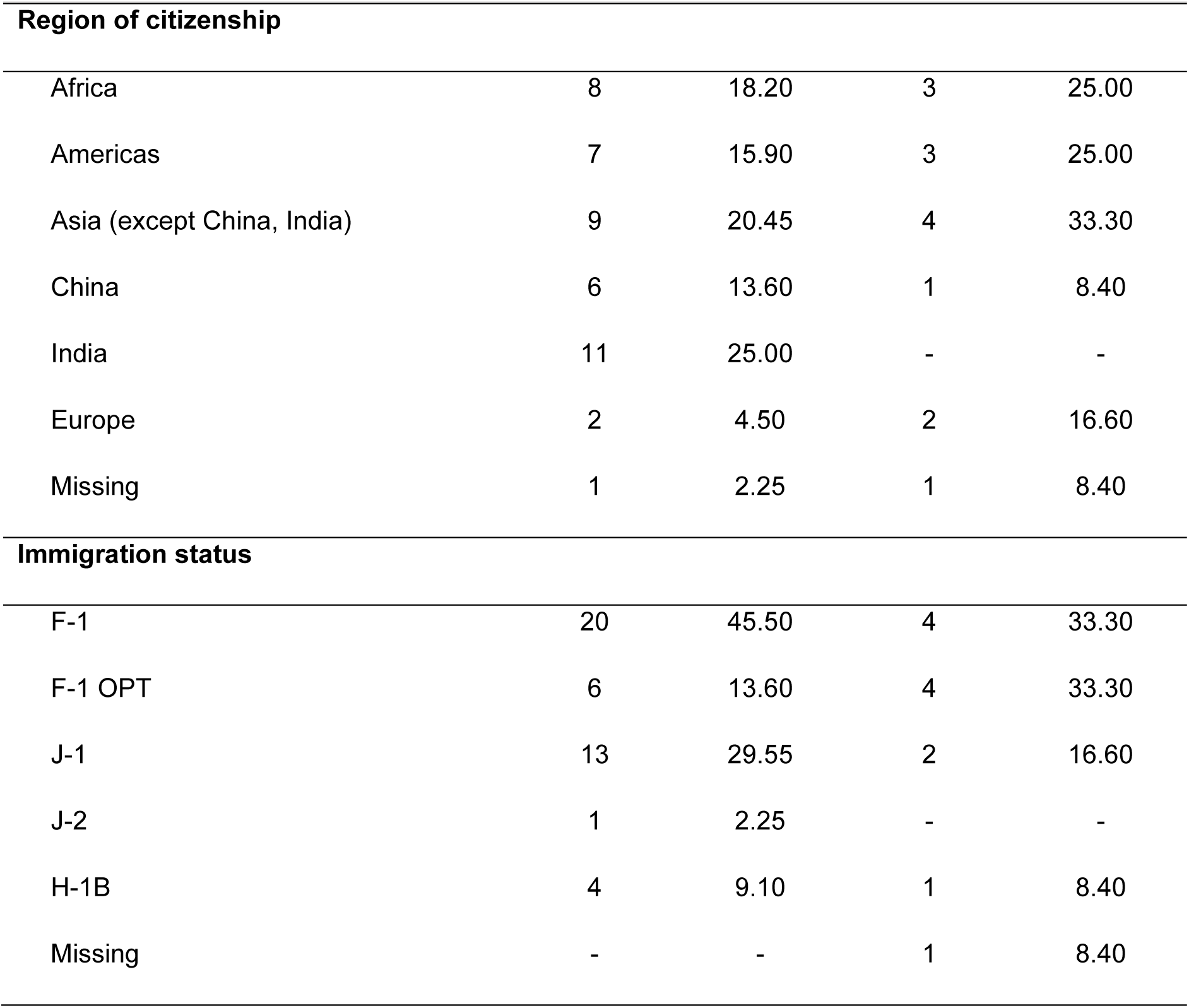
Demographics of Participants.

|  | Pre-program survey |  | Post-program survey |  |
| --- | --- | --- | --- | --- |
|  | n | % | n | % |
| Participants | 44 | 100.00 | 12 | 100.00 |
| **Training level** |  |  |  |  |
| Graduate student | 18 | 40.90 | 4 | 33.30 |
| Postdoctoral researcher | 23 | 52.30 | 7 | 58.30 |
| Others | 3 | 6.80 | - | - |
| Missing | - | - | 1 | 8.40 |
| **Location** |  |  |  |  |
| Drexel University | 22 | 50.00 | 6 | 50.0 |
| Washington University in St. Louis | 22 | 50.00 | 5 | 41.60 |
| Missing | - | - | 1 | 8.40 |
| **Gender identity** |  |  |  |  |
| Woman | 22 | 50.00 | 5 | 41.60 |
| Man | 21 | 47.75 | 5 | 41.60 |
| Prefer not to say | 1 | 2.25 | 1 | 8.40 |
| Missing | - | - | 1 | 8.40 |
| **Age range** |  |  |  |  |
| 18-25 | 4 | 9.10 | 1 | 8.40 |
| 26-35 | 32 | 72.70 | 7 | 58.30 |
| 36-45 | 8 | 18.20 | 3 | 25.00 |
| Missing | - | - | 1 | 8.40 |
|  | n | % | n | % |
| **Region of citizenship** |  |  |  |  |
| Africa | 8 | 18.20 | 3 | 25.00 |
| Americas | 7 | 15.90 | 3 | 25.00 |
| Asia (except China, India) | 9 | 20.45 | 4 | 33.30 |
| China | 6 | 13.60 | 1 | 8.40 |
| India | 11 | 25.00 | - | - |
| Europe | 2 | 4.50 | 2 | 16.60 |
| Missing | 1 | 2.25 | 1 | 8.40 |
| **Immigration status** |  |  |  |  |
| F-1 | 20 | 45.50 | 4 | 33.30 |
| F-1 OPT | 6 | 13.60 | 4 | 33.30 |
| J-1 | 13 | 29.55 | 2 | 16.60 |
| J-2 | 1 | 2.25 | - | - |
| H-1B | 4 | 9.10 | 1 | 8.40 |
| Missing | - | - | 1 | 8.40 |

Postdoctoral researchers were the largest fraction of participants (52% in the pre-program survey, 58% in the post-program survey). The distribution of participants by institution (Drexel University and Washington University in St. Louis) and by gender (self-identified as woman or man) was balanced. The largest proportion of participants (59% in the pre-program survey, 42% in the post-program survey) was from Asia, including China and India. Most participants reported holding F-1 immigration status, including those on post-graduation OPT (59% in the pre-program survey, 67% in the post-program survey). One participant from the post-program survey could not be linked to the pre-program survey and hence, demographics are missing for this person.

### 2.3. Data analysis

Data from the two surveys were organized by two members of the research team. Data from the polls were organized by one member of the research team, who was assisted by two graduate students who were not involved in the project otherwise. Additional help was used to collect post-program data that was qualitatively evaluated by one member of the research team. Whenever present, participants’ identifiable information was removed prior to analysis.

Pre- and post-program survey responses were linked via the participant’s chosen unique identifier. Data were analyzed in JASP (version 0.19) and Microsoft Excel (39,40). Results are represented either on the ordinal 5-point Likert scale or as the linear average of participants’ responses related to the introduced "Knowledge" and "Action" category (**Supplementary Source Data**). For statistical testing, we first tested the averaged, linear responses for "Knowledge" and "Action" against each other within a survey using the Wilcoxon signed-rank test. Secondly, we tested responses across surveys for participants who completed both surveys via a paired ANOVA and Tukey’s multiple comparison post hoc testing with *P*-value adjustment.

For the call-to-action polls, participants’ responses were accumulated from all polls (workshops #2-7) based on their names, and the analysis was conducted after the names were replaced with anonymizing identifiers. Rather than reporting on the completion of tasks on a weekly basis, we chose to report on the overall participation in these tasks across the entire program. A task was considered completed if the participant reported it as completed at any time point. The percentage of task completion was calculated based on the number of participants that responded to the poll question. For visualization, tasks were additionally categorized into three time investment groups: <15 minutes, 1-2 hours, and >3 hours.

Graphical representations of the data were generated with JASP (version 0.19), GraphPad Prism (version 10.4.1), and Adobe Illustrator (version 28.5) (39,41,42). Likert scale responses are shown on a 5-point scale (1-Strongly Disagree, 2-Disagree, 3-Neutral, 4-Agree, or 5-Strongly Agree) and on a 200% axis to visually separate disagreement (left side) from agreement (right side). Average Likert scale responses are presented as box plots showing 25% quartiles, median, and 95% confidence interval. Individual values are shown, and paired values are indicated via a line connection. Significance is shown as: *, <0.05; **, <0.01, ***, <0.001, and ****, <0.0001. Exact *P* values are given in the figure and their legends. Data references within the test contain the mean ± S.E.M., and other statistical values are summarized in the Supplemental Source Data file. Citations were created with Zotero (version 7.0.19) (43).

## 3. Results

We first evaluated participants’ self-assessment of their ability to complete the seven learning outcomes of the ECIP program (**Figure 1**). Pre-program survey data showed that participants had mixed responses related to the learning outcomes with a distribution throughout the 5-point Likert scale (**Figure 3A**). In contrast, post-program survey responses were clustered in the 4-Agree and 5-Strongly Agree categories. This data highlights the need for the program and underlines the growth opportunity for participants, while also indicating the substantial improvement across all learning outcomes after the program. Specifically, two learning outcomes showed the most progress: organizing an immigration portfolio (workshop #1) and selecting the right pathway to permanent residence (workshop #3). Additionally, the proportion of 3-Neutral responses decreased across all questions, suggesting that the program helped participants develop clearer, more confident perspectives on these learning outcomes. This trend was consistent among both graduate students and postdoctoral researchers (**Figure 3B, 3C**), indicating that the program is effective regardless of training level. These results provide evidence that the ECIP program achieved its intended learning outcomes.

**Figure 3.**
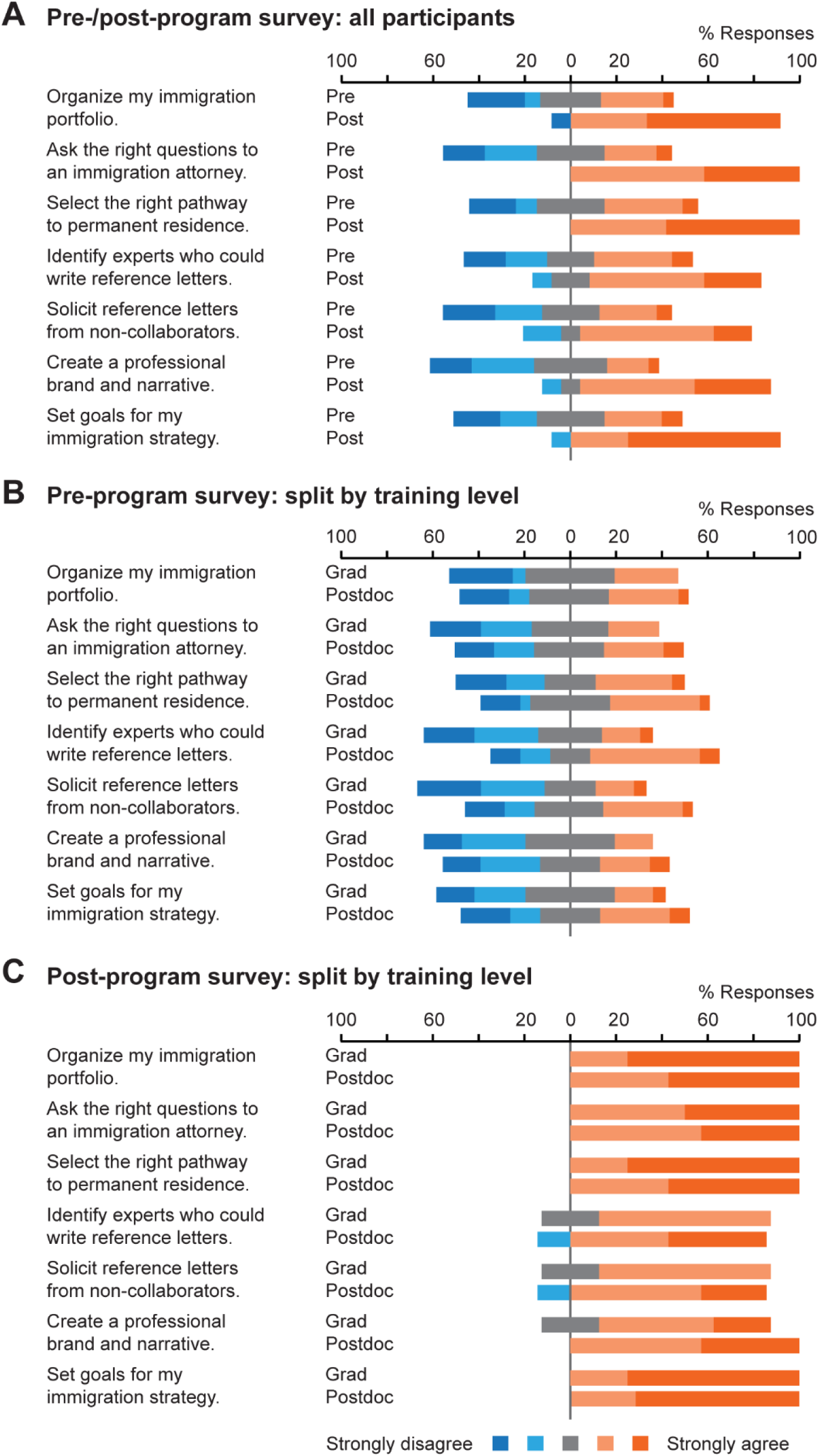
Participants’ self-assessment of the ECIP program learning outcomes before and after the program and within each training group. Participants’ pre- and post-program assessment of the seven workshops’ learning outcomes shown for (**A**) all participants in the pre-program versus post-program survey, (**B**) graduate students versus postdoctoral researchers in the pre-program survey, and (**C**) graduate students versus postdoctoral researchers in the post-program survey. Statistics: (A) Pre: pre-program survey (n=44). Post: post-program survey (n=12). (B) Grad: graduate student (n=18); Postdoc: postdoctoral researcher (n=23). (C) Grad: graduate student (n=4). Postdoc: postdoctoral researcher (n=7). Likert Scale: Dark blue: 1-Strongly Disagree; Light blue: 2-Disagree; Grey: 3-Neutral; Light orange: 4-Agree; Dark orange: 5-Strongly Agree.

As described in our study design, participants were asked to evaluate their level of understanding (“Knowledge”) of the program content. Across all participants, the proportion of responses in the 4-Agree and 5-Strongly Agree categories noticeably increased following program completion, especially in understanding how to prepare an immigration portfolio (workshop #1) and how to navigate permanent residence options (workshop # 3; **Figure 4A**). Topics that initially displayed lower levels of understanding (e.g., soliciting letters from unknown experts, building a professional brand) also showed improvement, revealing the program’s effectiveness in addressing previously less familiar content. We observed similar trends when splitting participants according to their training group (**Figure 4B, 4C**). These results demonstrate that the ECIP program had a positive impact on participants’ understanding of the intersection of career and immigration professional development topics.

**Figure 4.**
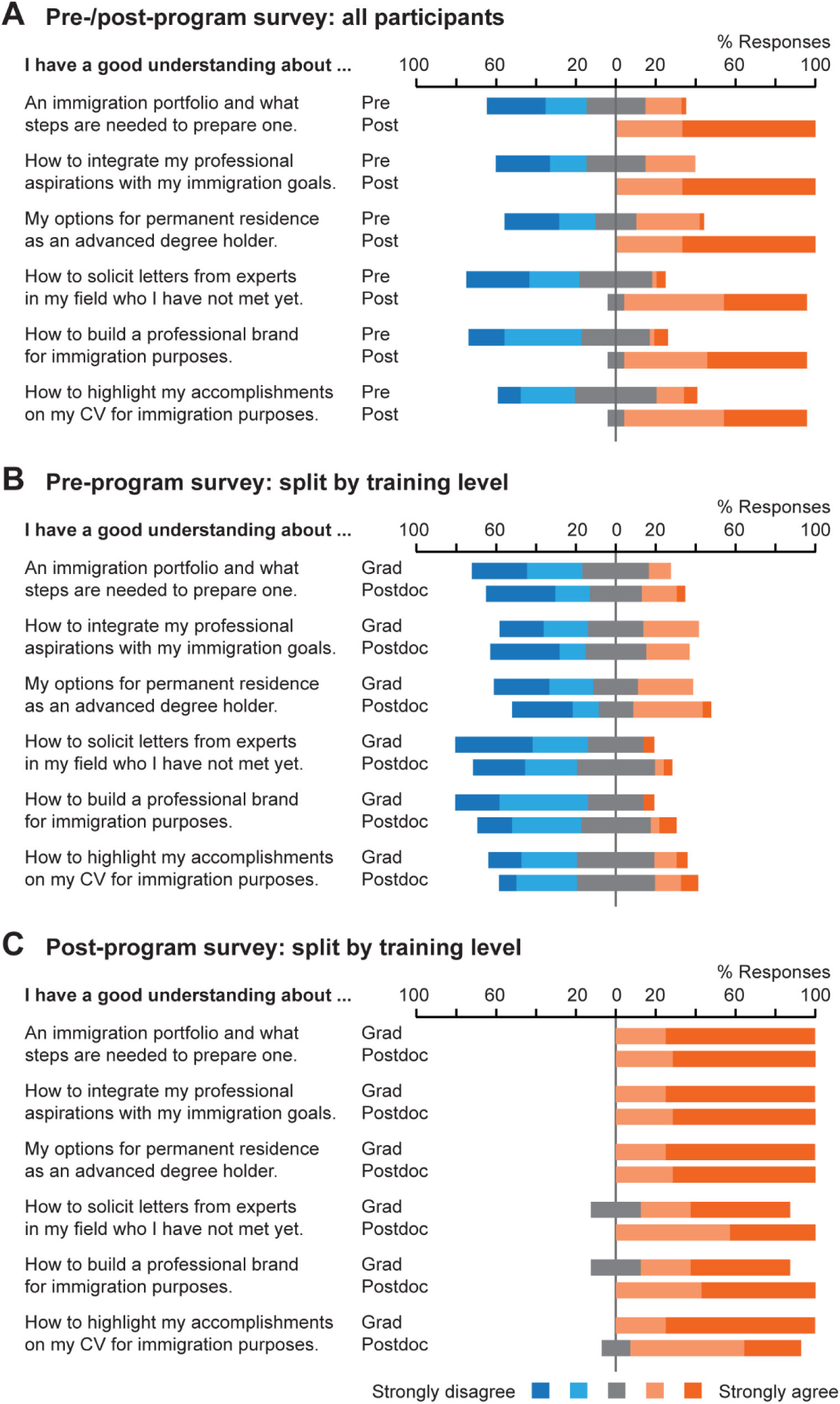
Responses for “Knowledge” category questions before and after the program and within each training group. Participants’ responses for a selected group of questions about their knowledge of the ECIP program content shown for (**A**) all participants in the pre-program versus post-program survey, (**B**) graduate students versus postdoctoral researchers in the pre- program survey, and (**C**) graduate students versus postdoctoral researchers in the post-program survey. Statistics: (A) Pre: pre-program survey (n=44). Post: post-program survey (n=12). (B) Grad: graduate student (n=18); Postdoc: postdoctoral researcher (n=23). (C) Grad: graduate student (n=4). Postdoc: postdoctoral researcher (n=7). Likert Scale: Dark blue: 1-Strongly Disagree; Light blue: 2-Disagree; Grey: 3-Neutral; Light orange: 4-Agree; Dark orange: 5-Strongly Agree.

In addition to their understanding of program topics that integrate career and immigration planning, participants were also asked about their confidence in taking related actions ("Action"). Comparison between these two quality groups in the pre-program survey revealed a significant difference in the response mean for “Knowledge” and “Action”, with an average of 2.45 ± 0.13 and 3.34 ± 0.15, respectively (Wilcoxon signed-rank test, W=26, ****, *P*<0.0001; **Figure 5A**). This difference suggests that participants were predisposed to take actions toward their career and immigration goals despite lacking the relevant knowledge. In the post-program survey, responses for “Knowledge” and “Action” clustered between the 4-Agree and 5-Strongly Agree categories. The average scores for these quality groups converged, reaching 4.48 ± 0.12 and 4.44 ± 0.10, respectively. Analogous results were observed when restricting the analysis to participants who completed both pre- and post-program surveys (paired ANOVA, *F*(2.054, 20.54)=36.95, ****, *P*<0.0001; **Figure 5B**). Examining the difference between “Knowledge” and “Action” within the graduate student and postdoctoral researcher group, we found the same robust difference in the pre-program survey and no difference in the post-program survey (**Figure 5C, 5D**). These quantitative results complemented our previous qualitative results in emphasizing the success and positive impact of the program.

**Figure 5.**
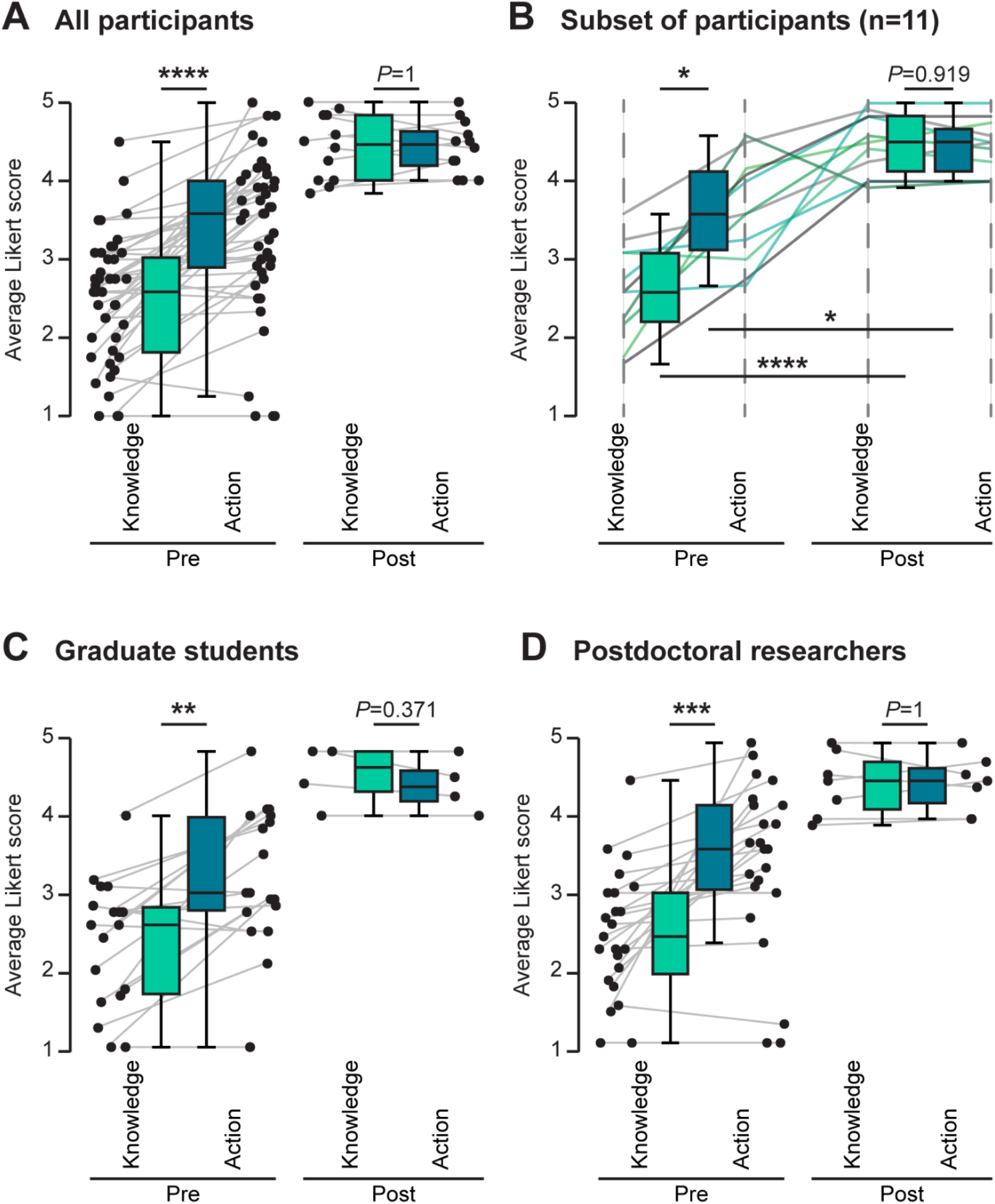
Alignment of “Knowledge” and “Action” category scores among participants after the ECIP program. Average Likert scoring per participant for Knowledge- and Action-type questions in the pre- and post-program survey. Individual paired measurements are shown for (**A**) all participants, (**B**) subset of participants who completed both surveys, and (**C**) graduate students and (**D**) postdoctoral researchers across pre- and post-program surveys. Statistics: (A) Wilcoxon signed-rank test. Pre: n=44, W=26, *P*<0.0001. Post: n=12, W=18, *P=*1.000. (B) Paired ANOVA (*F*(2.054, 20.54)=36.95) with Tukey’s post hoc testing: Pre: *P*=0.011; Post: *P*=0.919; Knowledge: *P*<0.0001; Action: *P*=0.022; n=11. (C) Wilcoxon signed-rank test. Pre: n=18, *P*=0.001. Post: n=4, *P*=0.371.(D) Wilcoxon signed-rank test. Pre: n=23, *P*=0.0002. Post: n=7, *P*=1.000. Median with upper/lower 25% quartile and 95% confidence intervals.

Polls at the beginning of workshops #2-7 asked about whether participants had completed the suggested call-to-action tasks, which allowed us to investigate their engagement with the ECIP program. Several of these tasks reached a completion rate of over 75% by the end of the program (**Figure 6A**), indicating success in engaging participants with the content and motivating them to work on their career and immigration goals. We found no bias in either the topic (Immigration Portfolio, Letters & CV, and Assistance) or the time investment for participants’ engagement with call-to-action tasks.

**Figure 6.**
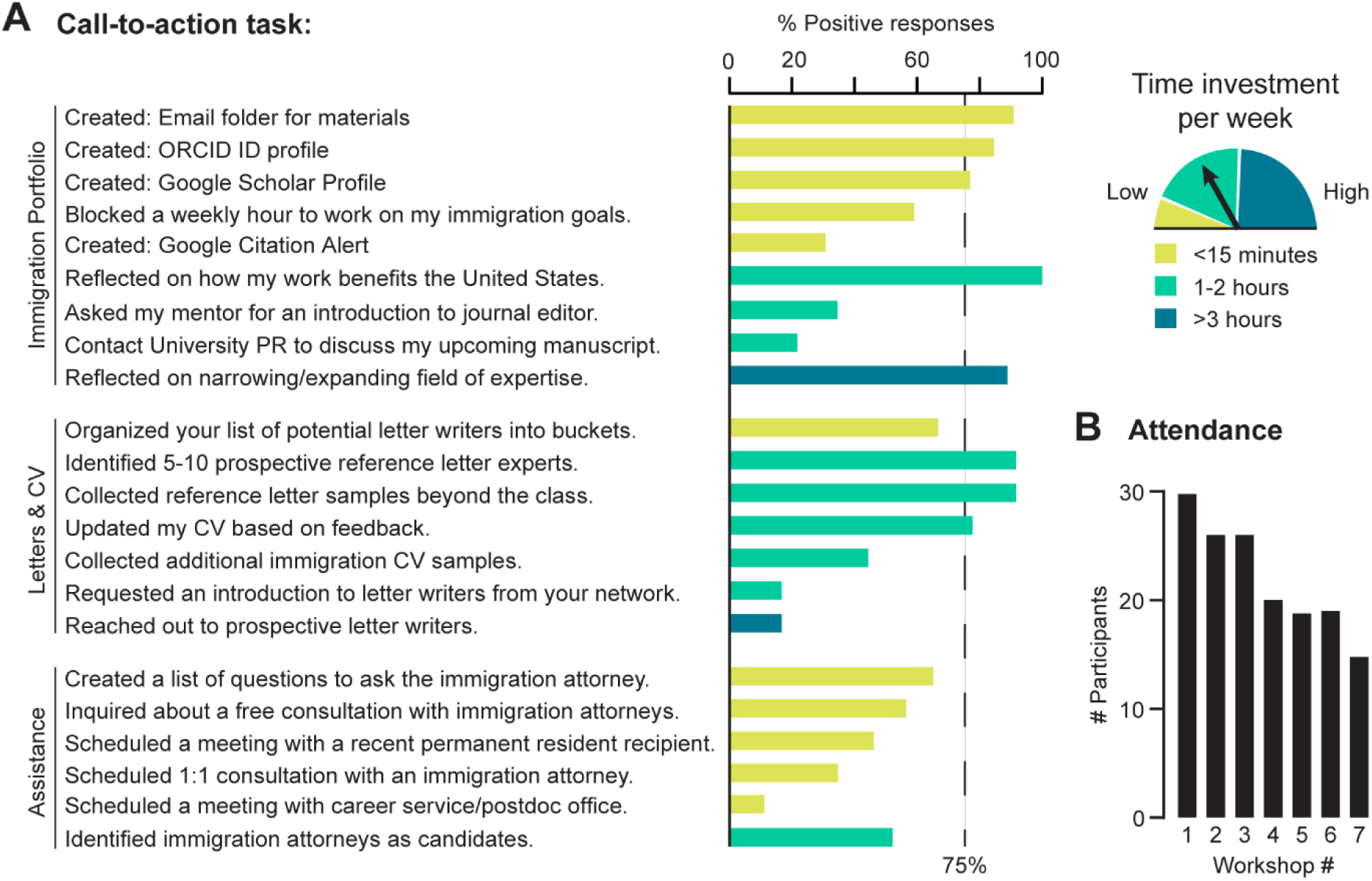
**Tasks completed by participants and attendance rate.** (**A**) Weekly call-to-action tasks reported as completed by participants (as % of responses) at the end of the program. Task topics are grouped as Immigration Portfolio, Letters & CV, and Assistance. Tasks are color-coded based on estimated time: Yellow: <15 minutes. Green: 1-2 hours. Blue: >3 hours. (**B**) Number of participants that attended workshops #1-7.

Additionally, we evaluated the participant attendance rate per workshop session (**Figure 6B**). The initial three workshops had the highest attendance rate (27.33 ± 1.33), followed by the workshops #4-6 (19.33 ± 0.33). Workshop #7 had the lowest participation, equivalent to approximately 50% of the initial attendance rate. These fluctuations could be explained by scheduling conflicts, program length, or participants’ perceived interest in the topic. For example, workshop #7 was considered of low interest by respondents of the post-program survey (**Supplementary Source Data**).

Evaluation responses within the post-program survey revealed overwhelmingly positive feedback both on the quality and the success of the ECIP program (**Figure 7A**). This feedback aligned with the program’s noted effectiveness to address critical knowledge gaps and boost confidence in taking immigration-related actions (**Figure 5**). Follow-up communications with eight participants who completed the program reaffirm these results, with four respondents indicating their active work on a permanent residence petition and several other replies regarding the positive outcomes and benefits of the program (**Figure 7B**). Participants reported a high return on investment, emphasizing the confidence they gained in their career and immigration planning, including learning relevant information and preparing their immigration portfolio. For example, they completed simple actions, such as saving documents (e.g., invitations, certificates, acknowledgments) for future use. They also described a new attitude to maximize activity outcomes, such as attending a conference not only for research purposes but also to keep in touch with experts who may eventually write recommendation letters for their immigration petition.

**Figure 7.**
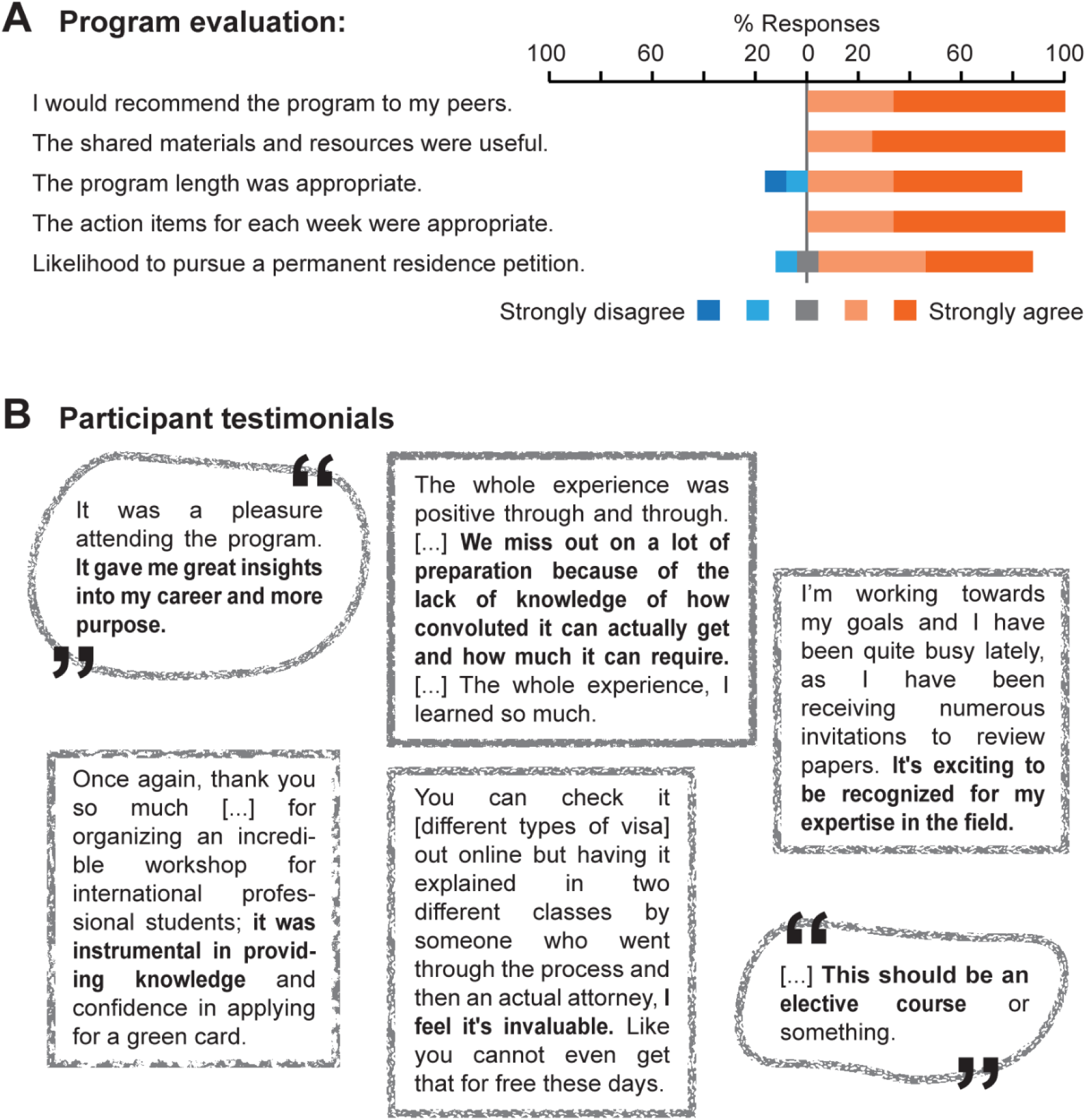
**Participant post-program evaluation and feedback after the program.** (**A**) Post-program survey responses related to the quality and satisfaction of participants after the program. Likert Scale: Dark blue: 1-Strongly Disagree; Light blue: 2-Disagree; Grey: 3-Neutral; Light orange: 4-Agree; Dark orange: 5-Strongly Agree. (**B**) Participant testimonials from follow-up communications at three and six months after the program.

## 4. Discussion

When aiming to launch a long-term career in the United States, international graduate students and postdoctoral researchers face a dual challenge: navigating a complex immigration system while making strategic career decisions in a competitive professional landscape. Many international scholars are unaware that their academic accomplishments can be used as evidence of eligibility for U.S. permanent residence. Despite their high representation in U.S. research institutions, these individuals often lack access to integrated resources that address how their immigration status influences their options for entering the U.S. workforce in academia, industry, or the public sector. The Effective Career and Immigration Planning for International Scholars (ECIP) program focuses on addressing this gap by offering a structured, educational intervention that aligns career development with immigration planning.

Results from this pilot study indicate that participation in the ECIP program led to improvements in participants’ self-assessed skills across all seven learning outcomes (**Figure 3A**) and their self-assessed knowledge regarding career and immigration decisions (**Figure 4A**). The most substantial gains were observed in participants’ understanding of how to prepare an immigration portfolio and navigate permanent residence options, which are central to the intersection of career and immigration planning. Likewise, the program significantly enhanced participants’ confidence to take action towards their career and immigration goals (**Figure 5A**). The aligned medians of "Knowledge" and "Action" responses in the post-program survey suggest that the ECIP program not only informed participants but also empowered them to take strategic steps in their career development.

Participant engagement was high throughout the program, particularly in completing call-to-action tasks. Weekly polls showed consistent participation, and most participants completed tasks during the program (**Figure 6A**). Attendance was strongest in the first three workshops, with a decline in later sessions (**Figure 6B**), which may reflect scheduling conflicts or a lower perceived interest. These findings suggest that the ECIP program successfully motivated participants to engage with the curriculum and to apply content to their personal immigration and career planning. Future program iterations could offer more flexible, condensed, or modular formats to accommodate varying schedules. Participants overwhelmingly rated the program as relevant, empowering, and of high quality. Post-program survey responses showed strong agreement with statements about the program’s usefulness, clarity, and applicability (**Figure 7A**). Additionally, follow-up communications revealed that several participants were actively taking steps to prepare their permanent residence petitions. Qualitative feedback emphasized the value of learning about immigration strategy in a structured, supportive environment, especially given the scarce opportunities to access resources like this program in current university settings (**Figure 7B**).

Our analysis affirmed that the ECIP program met a critical need and had a meaningful impact on participants’ professional development, addressing a gap in their training. By equipping international scholars with tools that allow them to make informed, strategic decisions about their long-term career plans in the United States, this study validated our hypothesis that structured guidance, linking academic accomplishments with immigration pathways, is highly beneficial for international scholars. The ECIP program brings essential attention to four deeply relevant areas in the experience of international scholars:

(1) Uncovering the hidden curriculum of career success;
(2) Shifting bias-driven assumptions to informed, intentional action;
(3) Enhancing the Individual Development Plan (IDP) framework to ensure relevance; and
(4) Supporting mental health.

### 4.1. Uncovering the hidden curriculum of career success

Through the ECIP program, participants learned how to leverage their academic and scholarly accomplishments with the goal of obtaining permanent residence—a necessary step for career success in academia, industry, and the public sector in the United States. By demystifying unspoken norms, timelines, and strategies, the ECIP program helps uncover the hidden curriculum at the intersection of career and immigration planning (28,29).

A consideration rarely addressed in career exploration is how to plan for the option of leaving the United States, regardless of citizenship or immigration status. While career pathways for doctoral recipients vary from country to country, a long-term career in any country relies on a robust professional network. The ECIP program offers participants the opportunity to reflect on strategies related to geography (workshop #1), regardless of whether they decide to return to their home countries, pursue career opportunities in another country, or remain in the United States.

As the ECIP program guides participants to undertake specific activities that support both their immigration and professional goals (e.g., skill building, networking, and document preparation), it empowers them to make proactive, strategic plans regarding their immigration portfolio. This approach is particularly important for scholars from countries with long permanent residence waiting times who must plan years in advance to remain competitive and compliant. In this sense, our results illustrate the crucial role of self-efficacy and contextual support in career decision-making (32,34). The success of the ECIP program emphasizes the vital importance of specialized career training for international scholars, preventing misconceptions and thus costly errors related to immigration.

### 4.2. Shifting bias-driven assumptions to informed, intentional action

Both graduate students and postdoctoral researchers in this study reported an initial bias to action, that is, a strong motivation to take immigration-related actions despite reporting a limited knowledge of the steps to do so (**Figure 5A**). This bias to action could affect as many as 73% of prospective international students, who would stay in the United States after graduation if the U.S. immigration structure were to be open to them (44). Without sufficient understanding about their career and immigration opportunities, a sense of urgency to act (likely driven by visa timelines and career pressures) can lead to poor decision-making and costly mistakes.

The perceived urgency to engage in actions underscores the need for timely, accessible, and targeted educational resources. The ECIP program successfully transformed participants’ motivation to act into informed motivation. The post-program survey results showed that both graduate students and postdoctoral researchers report alignment between "Knowledge" and confidence to engage in "Action" following completion of the ECIP program (**Figure 5B, 5C**). By effectively addressing their knowledge gap, the ECIP program enabled participants to act strategically to achieve their career and immigration goals.

### 4.3. Enhancing the IDP framework to ensure relevance

The IDP is a foundational evidence-based tool for career exploration and goal setting regardless of discipline or training stage (45). IDPs have been used in research education to support the career readiness of graduate students and postdoctoral researchers (46,47). The outcome and impact of the IDP process can be assessed through a set of instruments for each step of the process, namely self-assessment, exploration, goal setting, and decision making (48).

Given the interconnected nature of career and immigration planning for international graduate students and postdoctoral researchers, the ECIP program provides a vital opportunity to enhance the IDP framework in a novel way. We argue that the IDP framework can enhance its proven impact by incorporating immigration planning to identify activities that serve the dual purpose of advancing an international scholar’s career while strengthening their immigration case. This dual-focus approach is especially critical given the consequential outcomes of immigration petitions, which can have long-term effects for scholars and their families.

Integrating immigration education into the IDP framework empowers international scholars to strategize proactively and aligns their career and immigration goals towards an informed career trajectory. Broader adoption of this revised IDP format will also benefit the scientific community by helping to retain highly skilled talent. Having already invested time and resources in the United States, international scholars are likely to be well-positioned to contribute to the U.S. economic growth, the scientific workforce, and leading innovation, provided that their immigration pathways and career development are strategically aligned.

### 4.4. Supporting mental health

Although not directly measured in our study, qualitative feedback suggests that the ECIP program alleviated some participants’ stress and anxiety associated with immigration uncertainty. Early planning and peer support reduce the psychological toll of procrastination and last-minute visa crises, which impact mental health and productivity. Future iterations of the program should include validated mental health measures to assess the effect size of this variable.

## 5. Limitations

The study presents limitations that must be acknowledged to provide an adequate context for the reported observations. A significant limitation is the small sample size in both the pre-program survey (n=44) and the post-program survey (n=12), which may restrict generalizability. To increase the statistical power of our survey data, we included a paired analysis of data provided by participants who completed both surveys (n=11; **Figure 5B**). Findings from all participant survey responses (**Figure 5A**) are reinforced by this smaller, paired analysis, which demonstrated strong consistency between the responses of the larger group and those of the subset. Likewise, we could not analyze survey data by visa type, country of origin, or other demographic variables (**Table 1**) due to the small overall sample size. These variables may influence participants’ experiences and should be explored in future studies with a larger sample size to identify nuanced differences.

The presented analysis did not delve into the specific motivations and obstacles participants perceived regarding their immigration status. During workshop interactions, certain trainees voiced worries about maintaining their status, particularly due to timing issues associated with the permanent residence application process. This suggests that their challenges stemmed from the complexities of understanding correct timelines for their application rather than their ineligibility for permanent residence.

While the program was designed for U.S.-based scholars, its principle is broadly adaptable to international scholars in other countries. Specific adaptation would be necessary given that immigration systems vary widely.

Overall, while important insights were gained from this study, future program iterations will be critical in addressing these limitations and enhancing our understanding of the pervasive challenges advanced degree holders on temporary visas face in their academic and professional journeys.

## 6. Conclusion

The integration of career and immigration goals is essential for international graduate students and postdoctoral researchers. The presented ECIP program offers a scalable, evidence-based model for integrating career and immigration planning for international scholars. By aligning professional development with immigration preparedness through the immigration portfolio, the ECIP program addresses a critical gap in existing support structures. Program participants reported increased knowledge, confidence, and motivation to take strategic actions, which are essential for long-term success in the U.S. academic and professional landscape.

Given the high proportion of international scholars in U.S. research institutions, it is of national interest to retain this talent. Educational support like the ECIP program not only enables individual success but also contributes to the broader national goals of scientific innovation, economic growth, and workforce enrichment. Future iterations of this program should focus on: (1) expanding program access; (2) tailoring content to accommodate participants at varying stages of the permanent residence petition process; (3) evaluating the impact on mental health; and (4) tracking participants’ long-term career outcomes.

## Acknowledgments

This work was supported by the Burroughs Wellcome Fund Career Guidance for Trainees Award in 2023 (Grant #1074617) (38). Any opinions, findings, conclusions, and recommendations expressed in this article are those of the authors and do not necessarily reflect the views of the Burroughs Wellcome Fund. The funder played no role in the study design, data collection and analysis, decision to publish, or preparation of the manuscript. We thank Ms. Esther Kwarteng and Ms. Rachel Safo for their help with data organization for this study and Dr. Peter S. Myers for his help with post-program data collection. Editing assistance was provided by InPrint: A Scientific Communication Network at Washington University in St. Louis (49).

## Author Contribution

Conceptualization: NC, PC.

Data curation: NC, PC.

Formal analysis: CF.

Funding acquisition: NC, PC.

Investigation: CF, NC, PC.

Methodology: CF, NC, PC.

Project administration: NC, PC.

Supervision: NC, PC.

Visualization: CF.

Writing - original draft: CF, NC, PC.

Writing - review and editing: CF, NC, PC.

## Data Availability Statement

All relevant data is provided in the manuscript and the Supplementary Source Data.

## Abbreviations

ANOVA: Analysis of Variance
CV: Curriculum vitae
EB-1: Employment-Based Immigration: First Preference
EB-2: Employment-Based Immigration: Second Preference
ECIP: Effective Career and Immigration Planning for International Scholars
Grad: Graduate student
IDP: Individual Development Plan
OPT: Optional Practical Training
Post: Post-program survey
Postdoc: Postdoctoral researcher
Pre: Pre-program survey
R1: Research 1: Very High Spending and Doctorate Production
U.S.: United States

